# TDAG8 (GPR65) Inhibits Intestinal Inflammation in the DSS-Induced Experimental Colitis Mouse Model

**DOI:** 10.1101/496315

**Authors:** Edward J. Sanderlin, Swati Satturwar, Heng Hong, Kvin Lertpiriyapong, Mona Marie, Li V. Yang

## Abstract

T cell death-associated gene 8 (TDAG8, also known as GPR65) is a proton-sensing G protein-coupled receptor (GPCR) predominantly expressed in immune cells. Genome-wide association studies identify TDAG8 as a susceptibility candidate gene linked to several human inflammatory diseases including inflammatory bowel disease (IBD), asthma, spondyloarthritis, and multiple sclerosis. In this study, our results demonstrate that mice deficient of TDAG8 exhibited more severe inflammatory phenotypes than wild-type mice in a chronic dextran sulfate sodium (DSS)-induced colitis mouse model. Several disease parameters, such as diarrhea, colon shortening, fibrosis, histopathological score, and mesenteric lymph node enlargement were aggravated in TDAG8-null mice in comparison to wild-type mice treated with DSS. Increased leukocyte infiltration and myofibroblast expansion were observed in colonic tissues of DSS-treated TDAG8-null mice. These changes may represent a cellular basis of the observed exacerbation of intestinal inflammation and fibrosis in these mice. In line with high expression of TDAG8 in infiltrated leukocytes, real-time RT-PCR revealed that TDAG8 mRNA expression was increased in inflamed intestinal tissue samples of IBD patients when compared to normal intestinal tissues. Altogether, our data demonstrate that TDAG8 suppresses intestinal inflammation and fibrosis in the chronic DSS-induced colitis mouse model, suggesting potentiation of TDAG8 with agonists may have anti-inflammatory therapeutic effects in IBD.

## Introduction

Genome-wide association studies (GWAS) have identified numerous genetic risk loci for chronic inflammatory diseases. Large-scale GWAS studies have identified T cell death-associated gene 8 (TDAG8, also known as GPR65) as a susceptibility candidate gene for several human chronic inflammatory diseases such as multiple sclerosis, asthma, spondyloarthritis, and inflammatory bowel disease (IBD) (1, 12, 14, 17). A recent study demonstrates that TDAG8-deficient mice are more susceptible to bacteria-induced colitis and an IBD-associated TDAG8 genetic variant (I231L) confers reduced TDAG8 signaling activity as well as impaired lysosomal function (25). These data suggest TDAG8 could negatively regulate inflammation in certain diseases such as IBD.

TDAG8 was initially discovered as a gene up-regulated during T cell activation and apoptosis (5, 23). TDAG8 is highly expressed on leukocytes and leukocyte-rich tissues such as the spleen, lymph nodes, and thymus. Biochemically, TDAG8 can be activated by acidic extracellular pH through the protonation of several histidine residues on the receptor extracellular domains and transduce downstream signals through the G_s_/cAMP and G_12/13_/Rho pathways (13, 15, 16, 18, 19, 35, 39, 40, 44).

It has long been observed that the inflammatory loci can be more acidic than noninflamed tissues and that acidic pH can alter the function of inflammatory cells, vascular cells, and other stromal cells (4, 7–9, 18, 24, 33, 40, 41). The ways in which immune cells sense extracellular pH within inflamed microenvironments and subsequently alter their phenotypes have only recently been investigated. The role of TDAG8 activation by inflammation-associated acidosis has been investigated both *in vitro* and *in vivo*. Functionally, both pro- and anti inflammatory effects of TDAG8 have been described (1, 11, 13, 16, 22, 25, 32, 34, 35, 43). TDAG8 has been reported to impede pro-inflammatory profiles of primary murine macrophages, T cells, and microglia (13, 16, 29, 34). Investigation of TDAG8 in animal models of acute lung injury, arthritis, experimental allergic encephalomyelitis, myocardial infarction, and bacteria-induced colitis have indicated TDAG8 functions to inhibit inflammation in a variety of inflammatory diseases (25, 32, 35, 43).

As aberrant TDAG8 function is associated with IBD development and progression (17, 25), we sought to further characterize the role of TDAG8 in IBD. IBD is a broad term covering both Crohn’s disease (CrD) and ulcerative colitis (UC) (45). IBD is characterized by recurrent, aberrant inflammation within the intestinal tissue. These two disease forms are distinct, yet have overlapping clinical and histopathological features. The exact etiology is unknown, however, a complex interaction between immunologic, environmental, and genetic constituents is believed to contribute to the disease onset and progression. We utilized a chronic dextran sulfate sodium (DSS)-induced colitis mouse model to investigate the role of TDAG8 in experimental colitis. We observed that TDAG8 is protective against colonic inflammation and IBD associated complications. TDAG8 knockout (KO) mice treated with DSS had more severe clinical phenotypes such as body weight loss, fecal score, colon shortening, and mesenteric lymph node enlargement when compared to DSS-treated wild type (WT) mice. Histopathological analysis revealed that TDAG8 KO mice also had more severe histopathological features, intestinal inflammation, leukocyte infiltration, intestinal fibrosis, and isolated lymphoid follicles than WT mice.

## Materials and Methods

### 2.1 Dextran sulfate sodium (DSS)-induced chronic colitis mouse model

Experimentation was performed when male and female wild-type (WT) and TDAG8 knockout (KO) mice reached 9 weeks old. TDAG8 KO mice were generated as previously described (36) and backcrossed 9 generations into the C57BL/6 background. WT and TDAG8 KO mice were maintained under specific pathogen-free conditions and were free from Helicobacter, Citrobacter rodentium, and norovirus. Colitis was induced using 3% (w/v) colitis grade dextran sulfate sodium (DSS) with molecular weight 36,000-50,000, Lot# Q1408 (MP Biomedical, Solon, OH) within the drinking water of mice (41). Mice drank 3% DSS solution or water *ad libitum*. To cycle between moderate-severe inflammation, mice were given 3% DSS in autoclaved tap water or autoclaved tap water alone for 4 cycles. Each cycle constituted 5 days 3% DSS (severe) followed by 2 days of water (moderate). Following the fourth cycle, water was switched back to 3% DSS for 2 final days. Mouse body weight and clinical phenotype score were measured each day. All animal experiments were performed according to the randomized block experimental design. Additionally, all mouse experiments were approved by the Institutional Animal Care and Use Committee of East Carolina University, Greenville, North Carolina in accordance with the *Guide for the Care and Use of Laboratory Animals* (Office of Laboratory Animal Welfare, NIH).

### 2.2 Clinical phenotype scoring

Assessment of colitis severity was determined using the clinical parameters of body weight loss and fecal score (41). Each day stool was collected from mice and assessed for presence of blood and consistency. Fecal scoring system consisted of the following: 0= normal, dry, firm pellet; 1= formed soft pellet with negative hemoccult test, 2= formed soft pellet with positive hemoccult test; 3= formed soft pellet with visual blood; 4= liquid diarrhea with visual blood; 5= no colonic fecal content; bloody mucus upon necropsy. Presence of micro blood content was measured using the Hemoccult Single Slides screening test (Beckman Coulter, Brea, CA).

### 2.3 Tissue collection and processing

Upon terminal day of chronic colitis induction, mice were euthanized followed by necropsy. The gastrointestinal track was removed for analysis. Colon length was measured from the ileocecal junction to anus. Colon was then removed from cecum and the colon lumen was cleared of fecal content by washing with phosphate buffer saline (PBS) and then opened along the anti-mesenteric border. The colon tissue was then fixed with 10% buffered formalin and cut evenly into distal, mid, and proximal sections for histologic evaluation. Mesenteric lymph nodes were also collected and measured with a caliper. The largest mesenteric lymph node was collected for histological analysis and measurement. Mesenteric lymph node volume was calculated using the formula (length × width^2^) π/6. Lymph nodes were then fixed with 10% formalin for histological analysis.

### 2.4 Histopathological analysis

Five μm sections of distal, middle, and proximal colon tissue segments were stained with hematoxylin and eosin (H&E) for analysis. Sample identification was concealed during histopathological analysis for unbiased evaluation. Board certified medical pathologists (S.S. & H.H.) evaluated common colitis histopathological features including inflammation, crypt damage, edema, architectural distortion, and leukocyte infiltration in a blinded manner as previously described. Each parameter was scored and multiplied by a factor corresponding to total disease involvement. Scoring criteria and methodology were similar as previously reported, however, additional criteria including colonic fibrosis were added to this study. Colon segments were stained with picrosirius red and Masson’s Trichrome stains for fibrosis analysis and graded for pathological fibrosis as previously described with minor adaptations under the supervision of pathologists (6).

### 2.5 Immunohistochemistry

Colon tissues and mesenteric lymph nodes were embedded in paraffin and serial five μm sections were performed for immunohistochemical analysis as previously described (41). Briefly, antigen retrieval was performed of colon and mesenteric lymph node sections followed by endogenous peroxidase blocking. Tissue segments were blocked with normal serum and stained with anti-Green Florescence Protein (GFP) (Abcam, ab6673, Cambridge, MA), anti-F4/80 (Invitrogen, SP115, MA5-16363, Waltham, MA), anti-CD3 (Sigma, C7930, St. Louis, MO), and anti-αSMA (Abcam, ab5694, Cambridge, MA) primary antibodies. The IHC detection system anti-goat HRP-DAB Cell and Tissue staining kit (R&D Systems, Minneapolis, MN) or Superpicture 3^rd^ Gen IHC Detection system (Invitrogen, Waltham, MA) were used. Following addition of secondary antibody, DAB (3,3’ - diaminobenzidine) incubation was performed for HRP detection. Pictures were taken using the Zeiss Axio Imager A1 microscope.

### 2.6 Isolated lymphoid follicle quantification

Serial tissue sections were stained with H&E of the colon. Distal, middle, and proximal colon segments were scanned using a light microscope at 40× and 100× magnification in a blinded manner and isolated lymphoid follicles (ILFs) were counted. Colon segments were measured using a caliper for length in centimeters. Data are presented as ILFs/centimeter.

### 2.7 Leukocyte and myofibroblast quantification

Distal colon tissue segments were randomly selected (*n*=5-6) from WT-DSS and TDAG8 KO-DSS mice. Slides were stained for macrophage F4/80, myofibroblast α-SMA, and T cell CD3 markers, respectively. H&E slides were used to quantify neutrophils based on the characteristic polymorphonuclear morphology under the guidance of board certified pathologists. Pictures were taken (n=10-12) sequentially starting from the anus of the distal segment of the colon at 400× magnification. ImageJ software was utilized for counting of cells per high power field of view (FOV). The sample identification was concealed during the counting of cells for blind analysis.

### 2.8 Real-time RT-PCR

Real-time RT-PCR was performed as previously described using the TaqMan predesigned primer-probe sets for β-actin (Hs01060665_g1) and TDAG8 (GPR65) (Hs002692477_s1) (41). Crohn’s and colitis cDNA array was purchased from Origene Technologies (Catalog #CCRT102, Rockville, MD) to assess TDAG8 gene expression in human inflammatory bowel disease lesions compared to non-inflamed intestinal tissues. All sample information can be found at the vendor’s website and in previous report (41).

### 2.9 Statistical analysis

All statistical analysis was performed using GraphPad Prism software. The unpaired t-test or Mann-Whitney test was used to compare differences between two groups. When comparing three or more groups the one-way ANOVA was performed and followed by the post hoc Newman-Keuls test. *P* < 0.05 is considered statically significant.

## Results

### 3.1 Genetic deletion of TDAG8 exacerbates intestinal inflammation and fibrosis in the chronic DSS-induced colitis mouse model

To characterize the role of TDAG8 in chronic colitis, we utilized the chronic DSS-induced mouse model for the reliable induction of chronic intestinal inflammation (3, 21). During the experiment, clinical parameters were assessed such as body weight loss and fecal blood and diarrhea score. Both wild-type (WT)-DSS and TDAG8 knockout (KO)-DSS mice body weight loss was normalized to WT and TDAG8 untreated control mice. No body weight loss difference was observed between WT-DSS and TDAG8 KO-DSS during the first cycle, however, TDAG8 KO-DSS mice trended more body weight loss with most significant weight loss occurring at the end of cycle two and cycle three compared to WT-DSS mice (Fig. 1A). Fecal scores also indicated heightened severity of TDAG8 KO-DSS mice compared to WT-DSS mice (Fig. 1B). Both WT-DSS and TDAG8 KO-DSS mice reached an average fecal score of 3 by the end of the first cycle, however, TDAG8 KO-DSS mice maintained a more severe score during cycle two and cycle three compared to WT-DSS mice. Interestingly, during cycle four the TDAG8 KO-DSS mice partially recovered compared to WT-DSS. Upon the terminal point of the experiment, macroscopic disease indicators were evaluated such as mesenteric lymph node expansion and colon length shortening. Expansion of mesenteric lymph nodes (MLN) is a common parameter for colonic inflammation in the DSS-induced colitis model. Untreated control mice had MLN volumes approximately 5-6mm^3^ for WT and TDAG8 KO mice. MLN volume was significantly increased in DSS-treated mice as WT-DSS mice had an average volume of ^~^20mm^3^. Interestingly, TDAG8 KO-DSS MLN expansion was almost 2-fold higher than WT-DSS, indicating the inflammation of TDAG8 KO-DSS was more severe than WT-DSS. Finally, colon length was measured to assess the degree of shortening, which corresponds to heightened DSS-induced inflammation. WT-DSS mice had ^~^7% shortening compared to WT-untreated mice. TDAG8 KO-DSS mice, however, had more than 13% colon shortening. These data are not yet statistically significant given the current sample size.

**Fig. 1.**
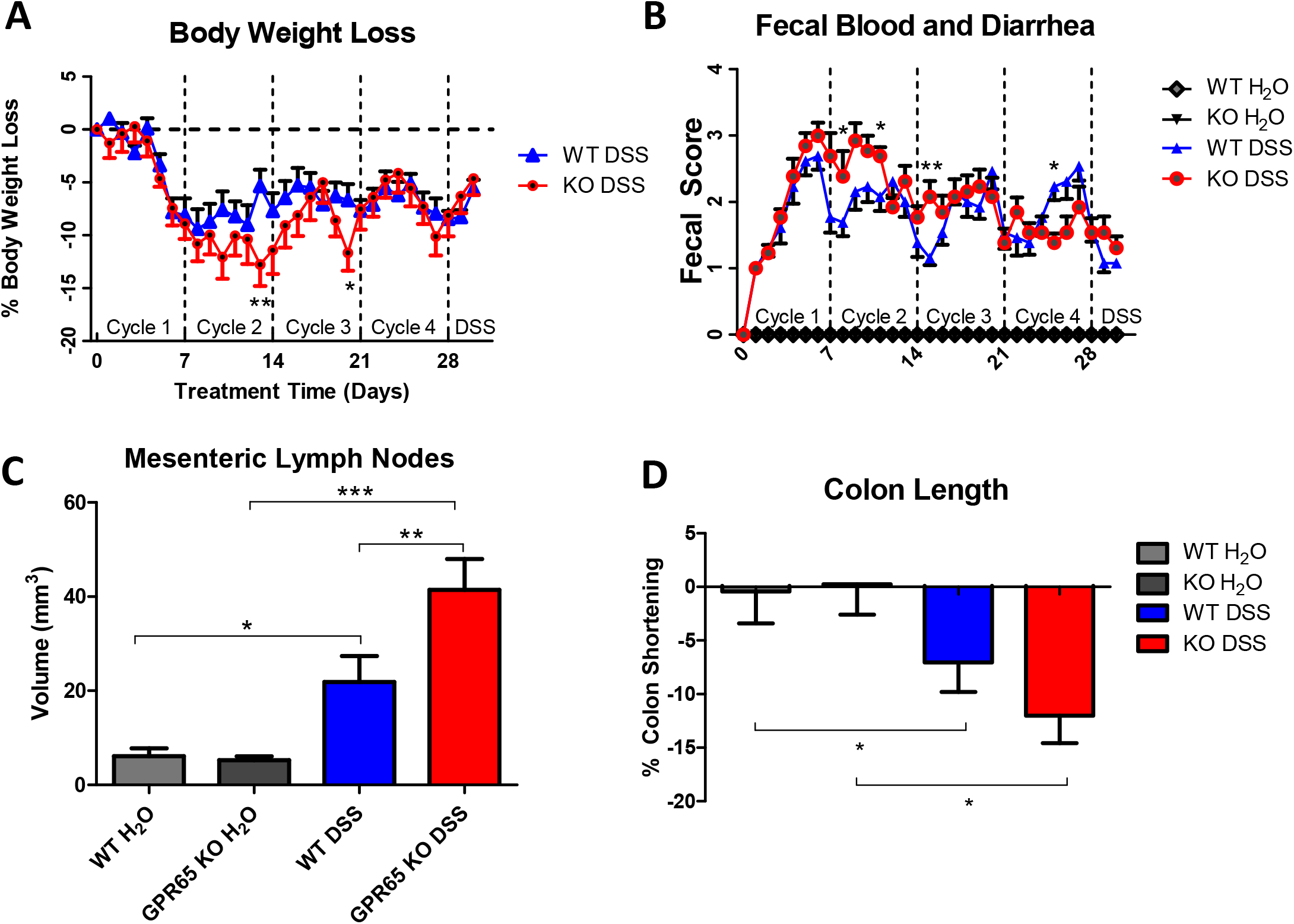
Disease indicators of chronic colitis induction in wild-type (WT) and TDAG8 knockout (KO) mice. The extent of DSS-induced inflammation was assessed in WT-DSS and TDAG8 KO-DSS mice compared to WT and TDAG8 KO control mice. TDAG8 KO-DSS mice presented elevated disease parameters compared to WT-DSS mice. Clinical phenotypes of intestinal inflammation such as (A) body weight loss and (B) fecal blood and diarrhea were assessed. Macroscopic disease indicators such as (C) mesenteric lymph node expansion and (D) colon shortening were also recorded. Data are presented as the mean ± SEM and statistical significance was determined using the unpaired t-test between WT-DSS and TDAG8 KO-DSS groups or one-way ANOVA followed by the Newman-Keuls Multiple Comparison test. WT control (N=10), WT-DSS (N=13), TDAG8 KO control (N=11), and TDAG8 KO-DSS (N=13) mice were used for experiments. (\**P* < 0.05, \*\**P* < 0.01, \*\*\**P* < 0.001).

To further assess the role of TDAG8 in intestinal inflammation, histopathological analysis was performed to quantify the degree of histological features of colitis between WT and TDAG8 KO mice. Distal, middle, and proximal segments of the colon were examined for common histopathological features of colitis, such as edema, crypt damage, architectural distortion, leukocyte infiltration, fibrosis, and inflammation. Untreated control mice had no colitis histopathological features (data not shown). We observed that the TDAG8 KO-DSS mice were more severe than WT-DSS mice in terms of total histopathology (Fig. 2A-D, I). Greatest histopathology was observed in the distal colon and is consistent with previous studies showing DSS model affects distal colon most severely.

**Fig. 2.**
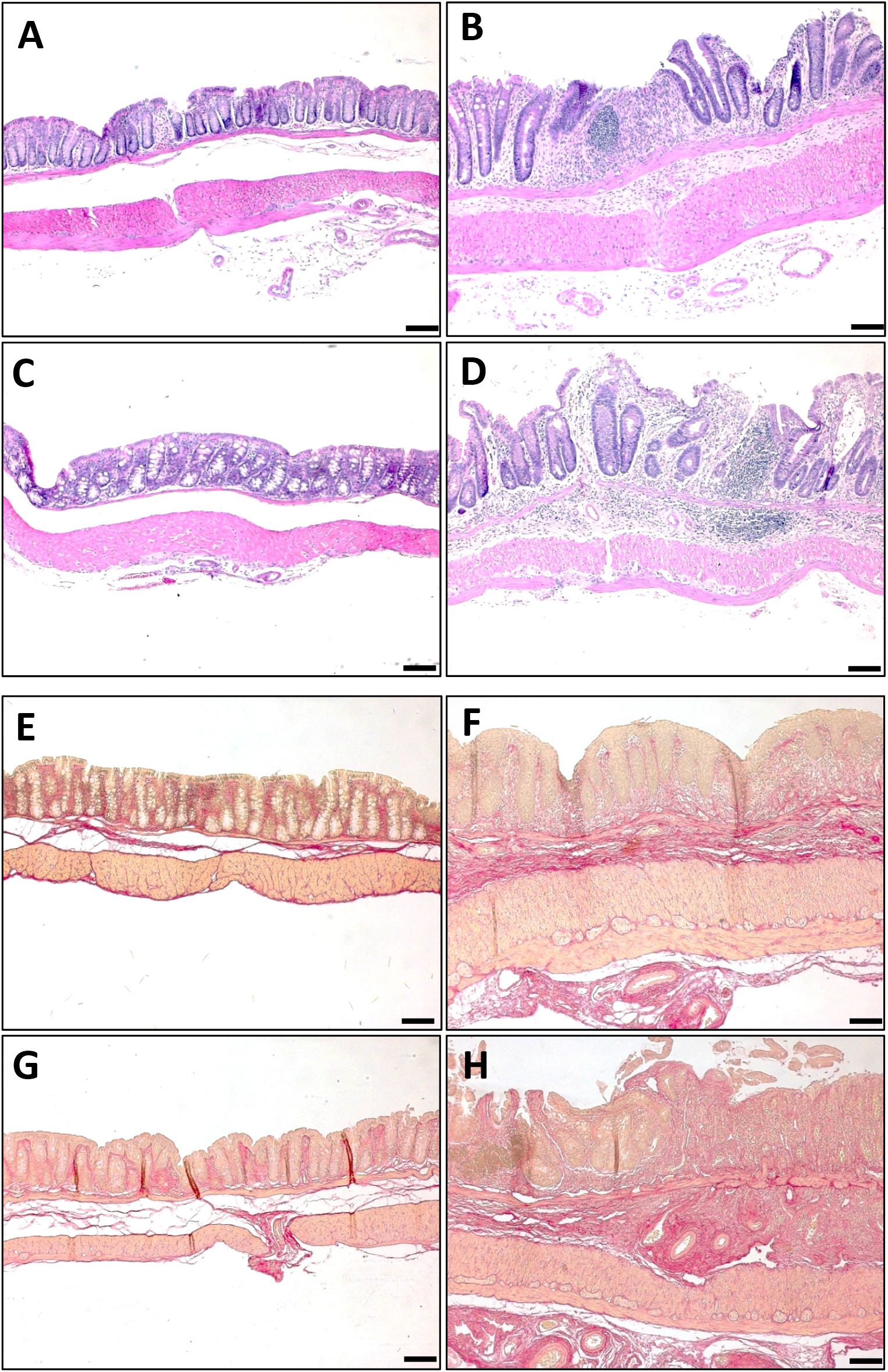
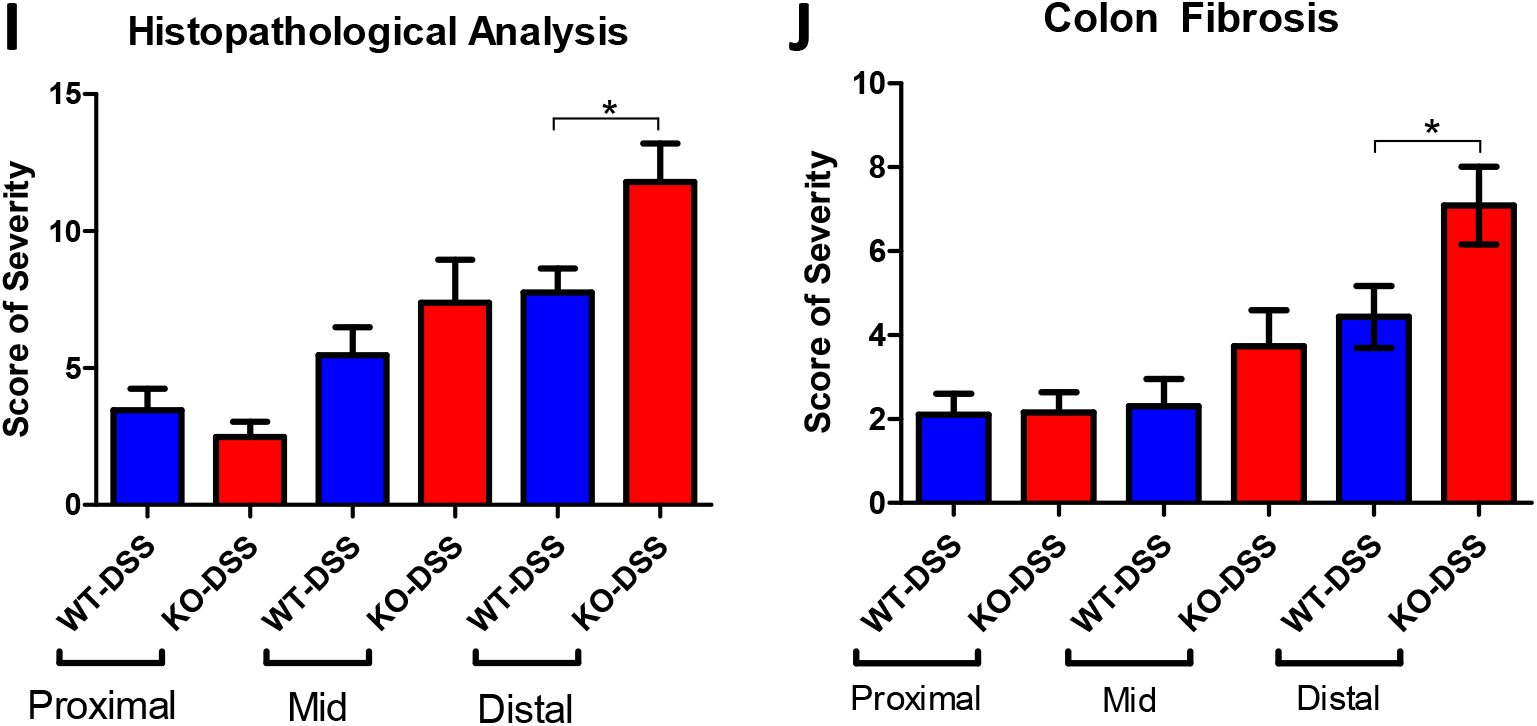
Histopathological analysis of proximal, middle, and distal colon. Characteristic histopathological features of colitis were assessed to further characterize the degree of intestinal inflammation. TDAG8 KO-DSS mice presented elevated disease parameters compared to WT-DSS mice. Representative H&E pictures were taken for (A) WT-control, (B) WT-DSS, (C) TDAG8 KO control, and (D) TDAG8 KO-DSS mice. Representative pictures of Picrosirius red stained tissue sections for fibrosis assessment were taken of (E) WT control, (F) WT-DSS, (G) TDAG8 KO control, and (H) TDAG8 KO-DSS mice. Graphical representation of (I) total histopathological parameters and (J) colonic fibrosis are presented. WT control (N=10), WT-DSS (N=13), TDAG8 KO control (N=11), and TDAG8 KO-DSS (N=13) mouse tissues were used for histopathological analysis. Scale bar is 100μm. Data are presented as the mean ± SEM and statistical significance was determined using the unpaired t-test between WT-DSS and TDAG8 KO-DSS groups. (\**P* < 0.05).

Fibrosis is another common pathological feature of IBD and can lead to life-threatening complications (27). We stained colon segments with picrosirius red and Masson’s trichrome for the assessment of colonic fibrosis after chronic intestinal inflammation (Fig. 2E-H). Distal, middle, and proximal colon segments were assessed for fibrotic development and scored accordingly for severity. Pathological fibrosis was assessed as the degree of increase compared to untreated controls. The greatest degree of fibrosis was found in both WT-DSS and TDAG8 KO-DSS distal colons with a gradual decrease in fibrosis towards proximal colon (Fig. 2J). We observed an almost 2-fold increase in fibrosis within distal colon of TDAG8 KO-DSS mice compared to WT-DSS mice. Severely fibrotic areas of the colon were characterized by increased collagen deposition within the lamina propria, muscularis mucosa, muscularis externa, and serosa when compared to control tissues (Fig. 2E-H, Supplementary Fig. 1).

Isolated lymphoid follicles (ILFs) are tertiary lymphoid tissues which can be induced within inflamed murine intestinal tissues. We have previously reported that ILFs are increased in intestinal tissues of mice in the DSS-induced acute colitis model when compared to untreated control mice (41). Here we demonstrate that ILF numbers are increased in areas of heightened colonic inflammation and are further increased in mice deficient of TDAG8 (Fig. 3). ILF numbers were counted in distal, middle, and proximal colon segments in untreated and DSS-treated WT and TDAG8 KO mice. ILFs were less prevalent in proximal segment when compared to middle and distal inflamed colon segments in both WT-DSS and TDAG8 KO-DSS mice (Fig. 3E). TDAG8 KO-DSS mice had increased ILFs/centimeter within each colon segment and subsequently full length colon when compared to WT-DSS (Fig. 3E-F). These data suggest TDAG8 prevents inflammation-associated ILF development.

**Fig. 3.**
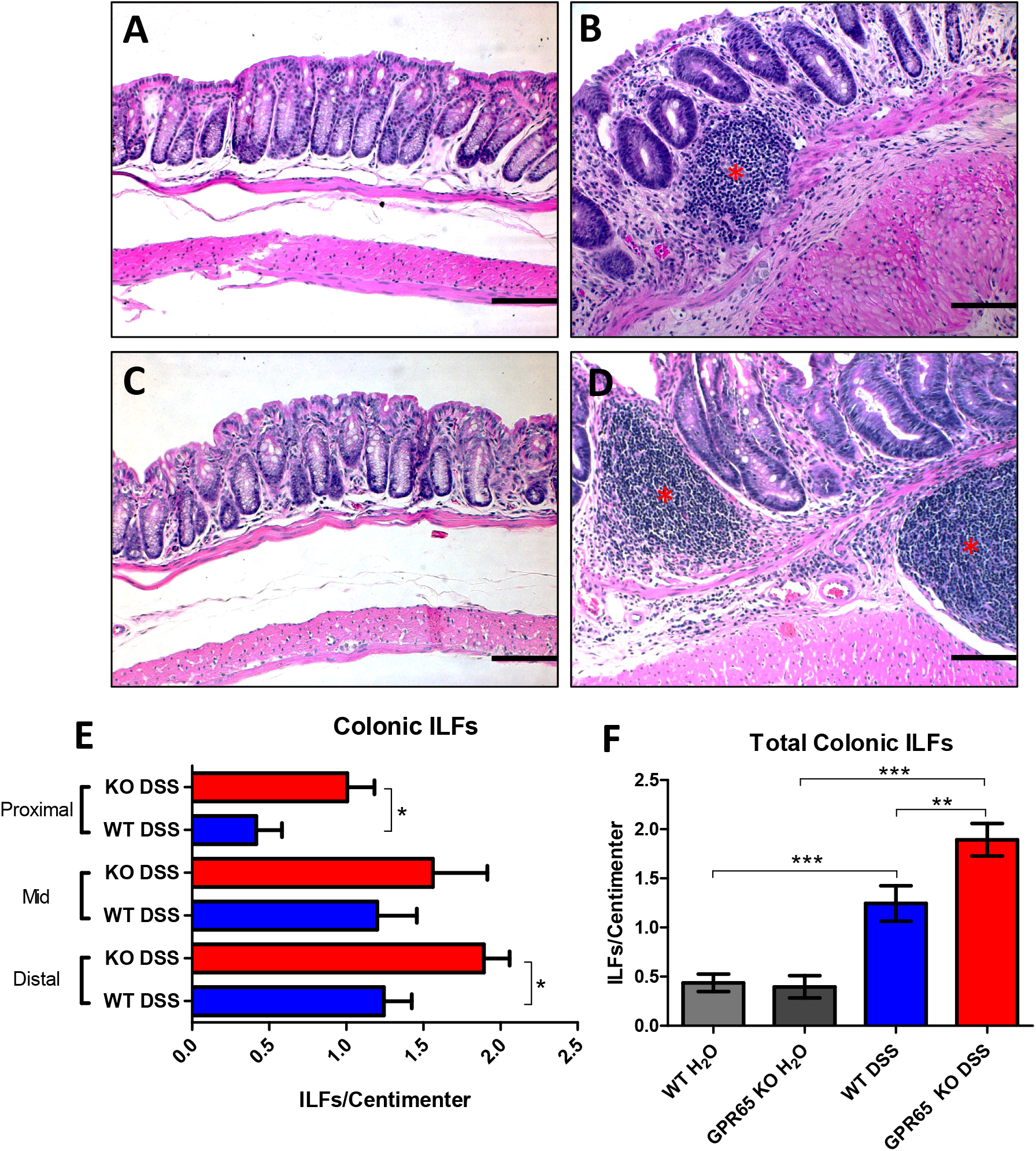
Isolated lymphoid follicle (ILF) quantification in proximal, middle, and distal colon segments. ILF numbers were assessed as an indicator of intestinal inflammation. ILF numbers were highest in distal colon with reduced numbers of ILFs in the proximal colon in DSS-treated mice. TDAG8 KO-DSS mice had a further increase in ILF numbers compared to WT-DSS mice. Representative pictures of ILFs in (A) WT-control, (B) WT-DSS, (C) TDAG8 KO-control, and (D) TDAG8 KO-DSS distal colon segments. Graphical representation of (E) ILF numbers in each segment of the colon and (F) combined full length colon. WT control (N=10), WT-DSS (N=13), TDAG8 KO control (N=11), and TDAG8 KO-DSS (N=13) mouse tissues were used for ILF quantification. Scale bar is 100μm. Data are presented as the mean ± SEM and statistical significance was determined using the unpaired t-test between WT-DSS and TDAG8 KO-DSS groups or one-way ANOVA followed by the Newman-Keuls Multiple Comparison test. (\**P* < 0.05, \*\**P* < 0.01, \*\*\**P* < 0.001).

### 3.2 Leukocyte infiltration and myofibroblast expansion are increased in the colon of DSS-treated TDAG8-null mice

To further investigate why TDAG8 KO-DSS mice had heightened fibrosis compared to WT-DSS mice, we examined the number of myofibroblasts within the inflamed and fibrotic distal colon mucosa. Myofibroblasts are one of several cellular constituents which can contribute to intestinal fibrosis (27). Normal intestinal myofibroblasts exist subjacent to the epithelium and regulate tissue repair, fibrosis, glandular secretion, and mucosal regeneration. α-SMA+ myofibroblasts numbers within noninflamed distal colon segments were quantified and compared to DSS-treated mouse colon segments (Fig. 4). There was a discernable increase in sub-epithelial mucosal myofibroblasts between WT-untreated and WT-DSS mice (^~^14/FOV vs. ^~^25/FOV) (Fig. 4G). Interestingly, TDAG8 KO mice had a further increase when TDAG8 KO untreated mice were compared to TDAG8 KO-DSS mice (^~^15/FOV vs. ^~^31/FOV). Very few myofibroblasts were observed within the submucosa (data not shown). Increased myofibroblast numbers could be detected within ulcerated areas of the colon when compared to non-ulcerated colon areas. Furthermore, during epithelial cell loss within ulcerated regions of the colon, myofibroblasts could be observed interspersing within disrupted epithelium for mucosal repair (Fig. 4C-D).

**Fig. 4.**
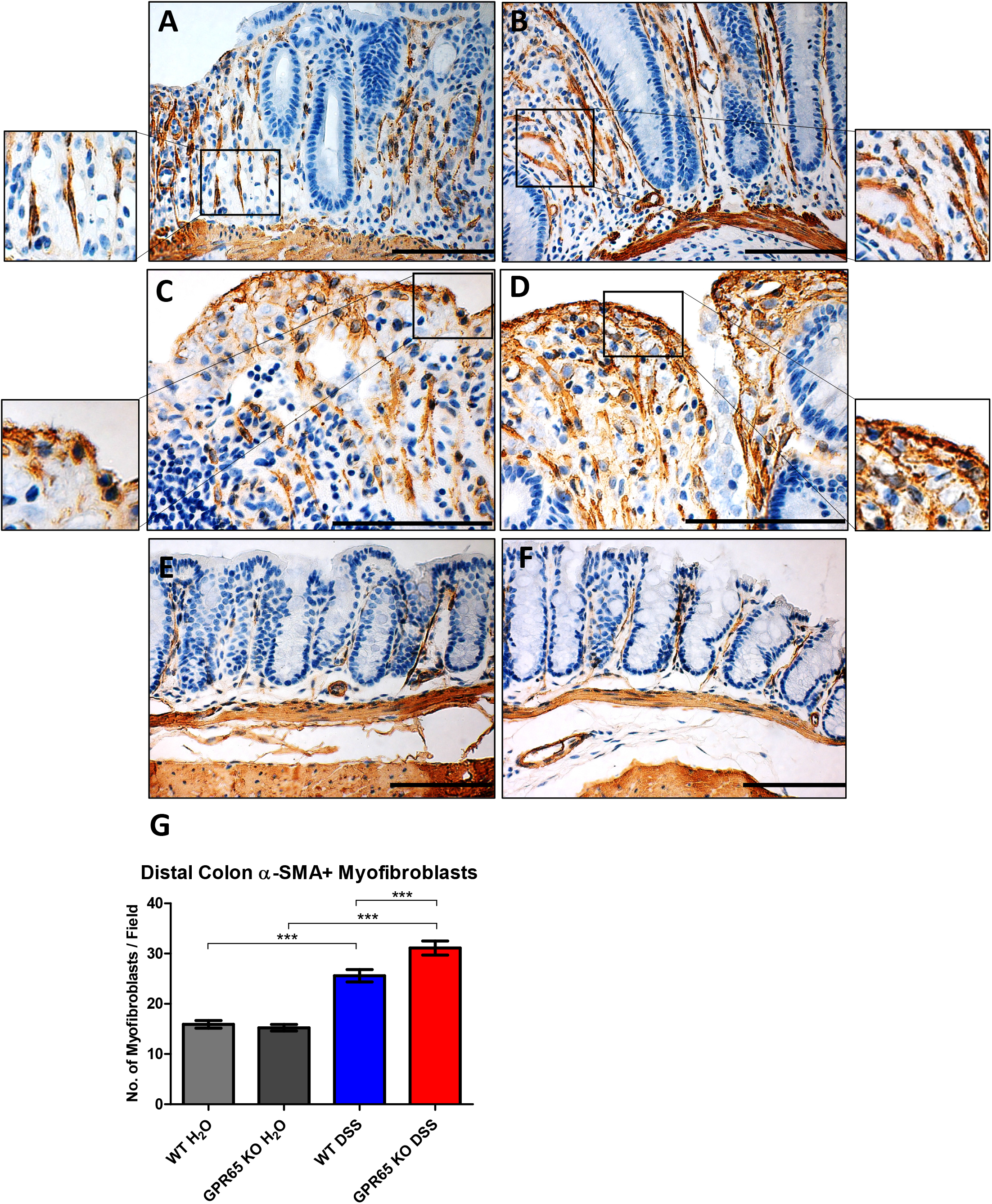
Myofibroblast expansion in distal colon mucosa. SMA^+^ myofibroblasts were quantified in distal colon as a cellular basis for increased colonic fibrosis. TDAG8 KO-DSS mice had increased myofibroblast numbers in distal colon compared to WT-DSS mice. Representative pictures of (A,C) WT-DSS, (B,D) TDAG8 KO-DSS, and (E) WT-control, and (F) TDAG8 KO-control. (G) Graphical representation of myofibroblast numbers in distal colon. WT-control (N=3), WT-DSS (N=6), TDAG8 KO-control (N=3), and TDAG8 KO-DSS (N=6). Scale bar is 100μm. Data are presented as the mean ± SEM and statistical significance was determined using the one-way ANOVA followed by the Newman-Keuls Multiple Comparison test. (\*\*\**P* < 0.001).

To further characterize colonic inflammation differences between WT-DSS and TDAG8 KO-DSS mice, we examined the populations of inflammatory cell infiltrates. The numbers of neutrophils, macrophages, and T cells within the distal colon were assessed. There was a significant increase in polymorphonuclear neutrophil, F4/80^+^ macrophage, and CD3^+^ T cell numbers between untreated and DSS-treated mice. We observed a 20-30% increase in leukocyte infiltrates within TDAG8 KO-DSS tissues when compared to WT-DSS colon tissues (Fig. 5).

**Fig. 5.**
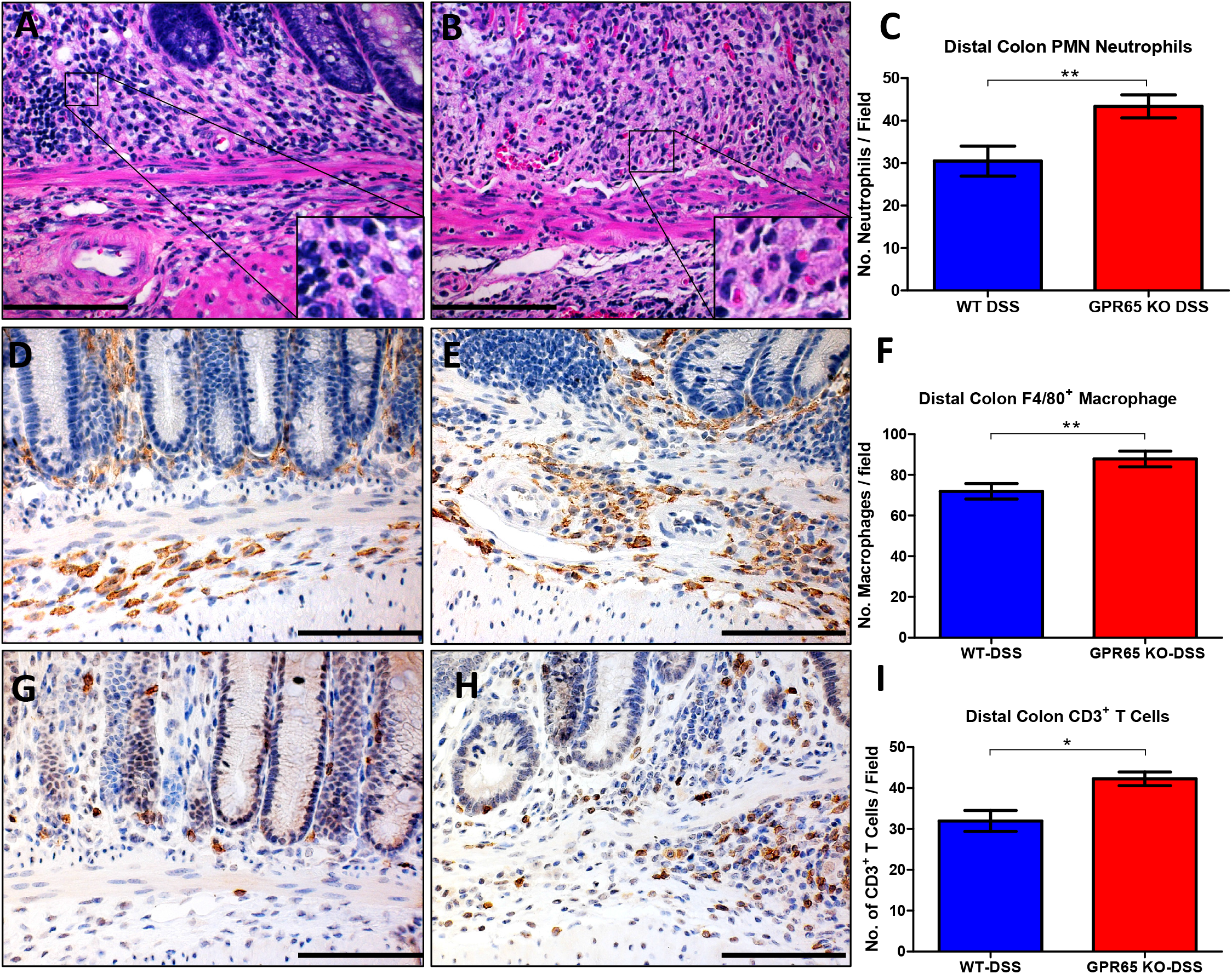
Leukocyte infiltrates in distal colon. Polymorphonuclear (PMN) neutrophils, F4/80^+^ macrophages, and CD3^+^ T cells were counted in the distal colon. TDAG8 KO-DSS mice had increased neutrophils, macrophages, and T cells in distal colon when compared to WT-DSS mice. (A) Representative pictures of WT-DSS and TDAG8 KO-DSS mouse (A-B) neutrophils, (D-E) macrophages, and (G-H) T cells, respectively. Graphical representation of (C) neutrophils, (F) macrophages, (I) and T cells. Scale bar is 100μm. Data are presented as the mean ± SEM and statistical significance was determined using the unpaired t-test between WT-DSS and TDAG8 KO-DSS groups. (\**P* < 0.05, \*\**P* <0.01).

### 3.3 TDAG8 gene expression in the mouse colon is predominantly detected in interstitial leukocytes

TDAG8 has been reported to be highly expressed in immune cells and leukocyte rich tissues such as the spleen, lymph nodes, and thymus (5, 23, 36). Increased TDAG8 expression can also be observed in tissues with higher baseline levels of resident leukocytes such as the lung and intestinal tissues. TDAG8 localization has not yet been investigated within the colon and mesenteric lymph nodes (MLN), most likely due to the lack of a reliable antibody. To investigate TDAG8 expression within the colon and MLN, we performed immunohistochemistry for green fluorescence protein (GFP) which functions as a surrogate marker for TDAG8 gene expression within TDAG8 KO mice (36). TDAG8 KO mice were generated by replacing the TDAG8 coding region with a promoterless internal ribosomal entry site-GFP cassette (36). Therefore, GFP expression is under the control of the endogenous TDAG8 promoter. GFP was detected in interstitial leukocytes within colon mucosa, transverse folds, and intestinal isolated lymphoid follicles (Fig. 6). Additionally, high GFP expression could be observed in MLN of untreated TDAG8 KO mice. Based on cellular morphology and localization, GFP expression within these colon leukocytes appears to be predominately macrophages, neutrophils, and lymphocytes. Within the MLN, high GFP expression was detected in histocytes and lymphocytes within the sinus regions, B cell follicles/germinal centers, and paracortical/interfollicular T cell zone. GFP expression was also assessed in DSS-treated TDAG8 KO mouse colon and MLN. A discernable increase of GFP positive leukocytes could be detected within the inflamed colon mucosa and transverse folds when compared to non-inflamed colon tissues. GFP could also be highly detected in ILFs and MLNs as observed in untreated mice. GFP expression is negative in both untreated and DSS-treated TDAG8 KO mouse epithelial cells and mesenchymal cells such as fibroblasts and smooth muscle cells. Additionally, endothelial cells are negative for GFP expression. There was also no GFP signal detected in any WT mouse tissues (Supplementary Fig. 2).

**Fig. 6.**
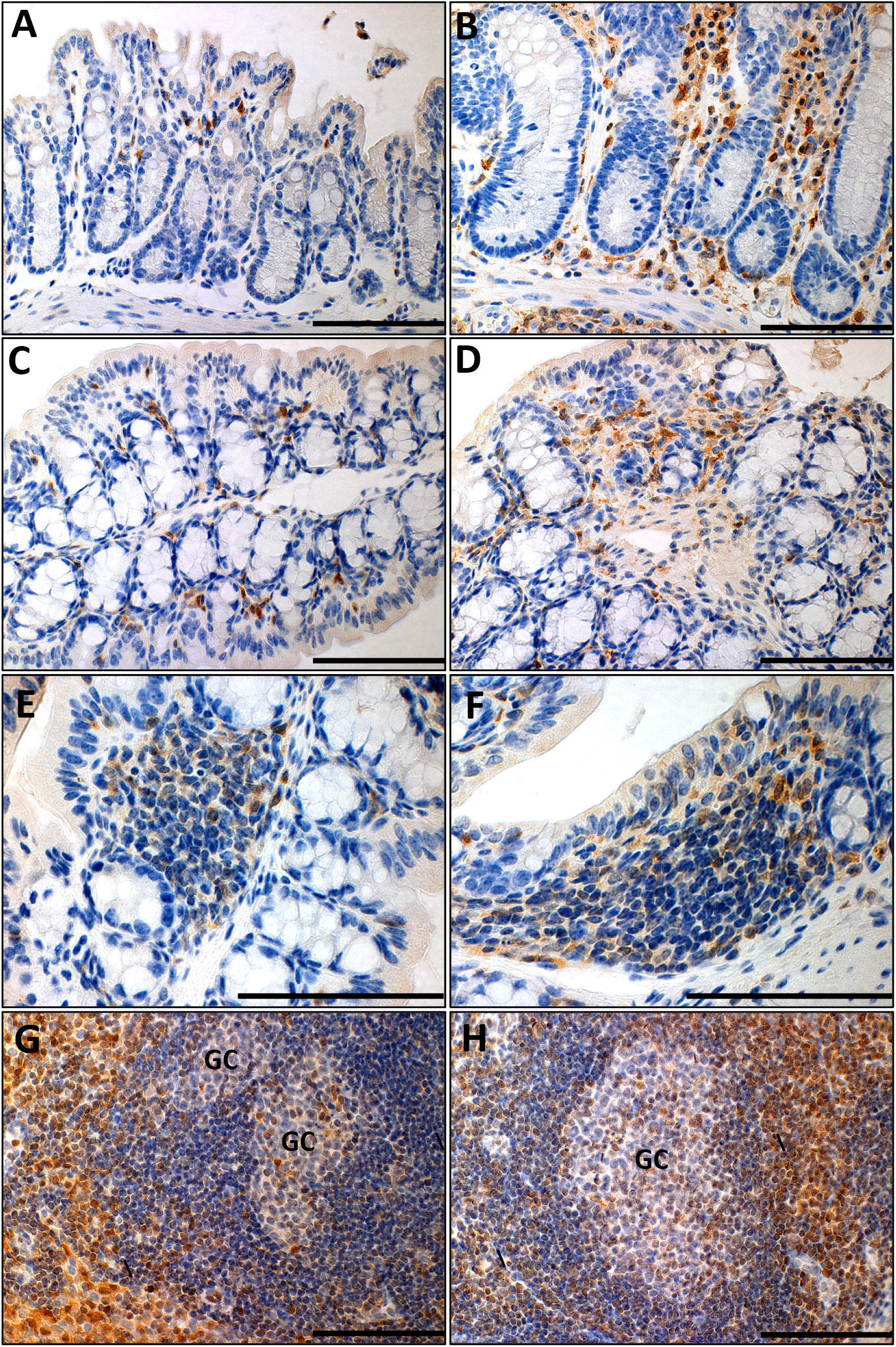
GFP signal in the intestine and intestinal associated lymphoid tissues. GFP knock-in signal serves as a surrogate marker for endogenous TDAG8 (GPR65) expression in TDAG8 KO mice. GFP signal could be detected in TDAG8 KO control (A) distal colon mucosa, (C) proximal colon transverse folds, (E) isolated lymphoid follicles (ILFs), and (G) mesenteric lymph nodes (MLNs). GFP signal could be detected in DSS-treated TDAG8 KO (B) intestinal mucosa, (D) transverse folds, (F) ILFs, and (H) MLNs. Based on cellular morphology and localization, GFP signal was observed in intestinal resident macrophages, lymphocytes, and neutrophils. GC: germinal center. Scale bar is 100μm.

### 3.4 TDAG8 gene expression is increased in inflamed intestinal tissues of IBD patients

TDAG8 gene expression was assessed in intestinal tissues from patients with ulcerative colitis (UC) and Crohn’s disease (CrD) in comparison to normal intestinal tissues. A cDNA array including 7 non-inflamed intestinal tissue samples, 14 CrD tissue samples, and 26 UC tissue samples was utilized. We observed a ^~^4-fold increase of TDAG8 gene expression in UC samples and a ^~^6-fold increase in CrD samples compared to normal intestinal samples (Fig. 7).

**Fig. 7.**
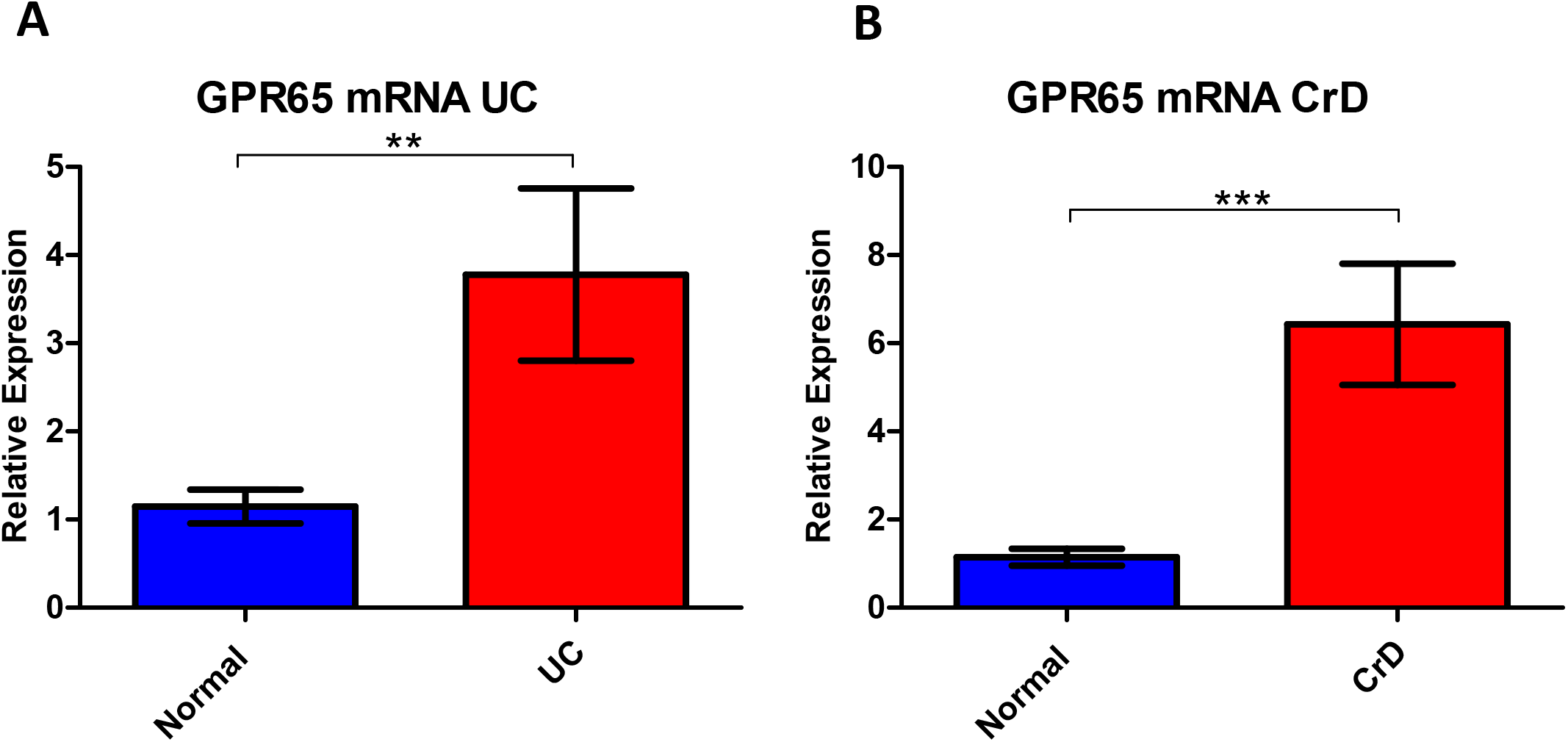
TDAG8 (GPR65) gene expression in human ulcerative colitis and Crohn’s disease. TDAG8 (GPR65) mRNA is increased in ulcerative colitis and Crohn’s disease compared to non-inflamed intestinal tissues. (A) TDAG8 (GPR65) gene expression in ulcerative colitis and (B) Crohn’s disease. Data are presented as the mean ± SEM and statistical significance was determined using the Mann-Whitney test between control and diseased intestinal tissues. (\*\**P* < 0.01, \*\*\**P* < 0.001).

## Discussion

In our study, we investigated the functional role of TDAG8 in a mouse model of chronic DSS-induced experimental colitis. Our results indicated that TDAG8 provides a protective role in reducing intestinal inflammation in the DSS-induced chronic colitis mouse model. TDAG8 deficiency in mice aggravated body weight loss, fecal scores, mesenteric lymph node expansion, and colon shortening in comparison to WT mice. Additionally, TDAG8 absence resulted in exacerbated histopathological features of IBD such as inflammation, edema, leukocyte infiltration, and fibrosis. Isolated lymphoid follicles (ILFs) were also expanded in TDAG8 KO colon tissues compared to WT. TDAG8 was also shown to be expressed in the colon infiltrated leukocytes and likely impedes inflammation through inhibition of immune cell inflammatory programs once activated by acidic pH in the inflamed intestinal loci. These results are supportive of previous studies demonstrating an anti-inflammatory role of TDAG8 in a diverse set of diseases (25, 32, 35, 43), including chronic intestinal inflammation and provide new insights into the role of TDAG8 in colitis.

The inflamed intestinal mucosa is characterized by leukocyte infiltration, and resulting tissue damage due to unresolved, chronic inflammation. Chronic inflammation can lead to acidic tissue microenvironments owing to increased metabolic byproducts of infiltrated immune cells and further alter the inflammatory response. Numerous studies have shown that local tissue pH below 7.0, and sometimes even below 6.0, is detected in inflammatory conditions, tumors, and ischemic tissues (9, 18, 24, 33, 40). Additionally, several reports indicate that patients with IBD have a more acidic colon when compared to control patients with pH ranging from less than pH 5 to slightly below pH 7.4 (9, 33). Therefore, acidosis and inflammation often co-exist in inflamed intestinal tissues. Mesenchymal cells and leukocytes will function within these acidic, inflamed microenvironments and either potentiate or inhibit local inflammation. Wound-healing mechanisms will also occur within acidic and inflamed intestinal lesions. pH-sensing is critical for cells to sense alterations in environmental proton gradients to maintain proper cellular functions. These microenvironmental conditions within inflamed tissues present a role for the pH-sensing GPCR TDAG8. TDAG8 can sense alterations in local pH by leukocytes and subsequently alter immune cell functions (13, 16, 22, 25, 32, 34, 43). In our current study, we provide evidence that genetic deletion of TDAG8 exacerbates intestinal inflammation in a chemically induced colitis mouse model. Further studies are warranted to assess alterations in local interstitial pH within inflamed lesions of the gut as well as the effects of tissue associated acidosis on both TDAG8 expression and function of immune cells.

Previous studies have characterized TDAG8 as a functional pH-sensor expressed predominately within leukocytes and can negatively regulate the inflammatory response of immune cells (13, 16, 29, 34). Furthermore, downstream effectors of the TDAG8-coupled Gα_s_/cAMP have demonstrated anti-inflammatory roles in a diverse set of processes (30). Gα_s_/cAMP/PKA/CREB pathway has been shown to reduce granulocyte, macrophage, and monocyte inflammatory programs (37). Additionally, cAMP can reduce dendritic cell function in lymph nodes, T cell activation, and can increase T regulatory cell activity. These data are consistent with reports that TDAG8 activation can inhibit inflammatory profiles in macrophages, microglia, neutrophils, and T cells (13, 16, 29, 31, 34). Additionally, the role of TDAG8 was investigated in immune-mediated murine disease models such as arthritis, lipopolysaccharide (LPS)-induced acute lung injury, myocardial infraction, ischemic stroke, and bacterial-induced colitis (25, 28, 32, 35, 43). However, there are also some studies suggesting TDAG8 expression in eosinophils promotes inflammation through increasing eosinophil viability in an asthma mouse model (22). Furthermore, recent studies found TDAG8 is a regulator for Th17 pathogenicity and increases the severity in the experimental autoimmune encephalomyelitis (EAE) mouse model as well as increases GM-CSF production in CD4 T cells (1, 11). Pertaining to Th17 cell pathogenicity, reports have provided evidence for both protective and pathogenic roles in the context of intestinal inflammation (10). In addition to these *in vitro* and *in vivo* animal studies, recent genome wide association studies (GWAS) have identified small nucleotide polymorphisms (SNPs) of TDAG8 associated with several human inflammatory-mediated disease states such as multiple sclerosis, asthma, heparin-induced thrombocytopenia, spondyloarthritis, and IBD (1, 12, 14, 17, 20). As previously mentioned, one group investigated an IBD-associated TDAG8 genetic variant (I231L) within a bacteria-induced colitis mouse model and found that this TDAG8 gene variant confers reduced TDAG8 activity as well as impaired lysosomal function (25). Additionally, a recent study found TDAG8 has protective effects in intestinal inflammation mouse models (42). In Our study focuses on the functional role of TDAG8 in the regulation of inflammation in a chemically induced chronic colitis mouse model and further provides evidence that TDAG8 functions to inhibit inflammation in colitis. We demonstrated that TDAG8 is expressed in infiltrated leukocytes within the colon of inflamed intestinal tissues using GFP as a surrogate marker in TDAG8 KO mice. GFP-positive leukocytes were predominately macrophages, neutrophils, and lymphocytes based on cellular morphology in comparison to F4/80 and CD3 immunostaining. There is a discernable increase in GFP positive leukocytes in the DSS treated mouse colon tissues compared to the untreated tissues indicating TDAG8 expression is increased in inflamed tissues compared to non-inflamed intestinal tissues. We then assessed TDAG8 gene expression in human colitis and Crohn’s intestinal lesions compared to non-inflamed intestinal tissues. We observed that TDAG8 is increased by more than 4-fold in inflamed intestinal tissues compared to control. It is likely the increased expression of TDAG8 in IBD intestinal samples is due to the increase of infiltrated leukocytes, which have high endogenous TDAG8 expression.

We also found that pathological fibrosis was increased in TDAG8 KO-DSS mice compared to WT-DSS mice. Fibrosis is a serious consequence of recurrent intestinal inflammation and can lead to complications such as intestinal strictures and obstruction (26, 27, 38). Collagen can be produced by several cellular constituents within the intestinal tissues. Some such cells include fibroblasts, sub-epithelial myofibroblasts, smooth muscle cells, and pericytes. Additionally, fibroblasts, smooth muscle cells, fibrocytes, endothelial cells, pericytes can undergo epithelial/endothelial-mesenchymal transition into myofibroblasts for wound healing functions (26, 27, 38). Myofibroblasts are described as a major contributor of pathological extracellular matrix deposition within the inflamed intestine (26, 27, 38). As such, we quantified the number of SMA^+^ myofibroblasts in the mucosa and observed TDAG8 KO mice had more myofibroblasts than WT mice, supporting the observed increased fibrotic deposition in the DSS-treated TDAG8 KO colon. It remains to be determined how TDAG8 regulates fibrosis in the chronic colitis model. Interestingly, however, a recent study demonstrates that TDAG8 regulates macrophage CCL20 expression, γδT cell infiltration, and fibrosis in a myocardial infarction mouse model (32).

Altogether, our results provide further support for an anti-inflammatory role of TDAG8 in colitis and present TDAG8 as a potential target for therapeutic intervention. Currently, IBD treatment options are limited and predominately consist of steroids, anti-TNFα monoclonal antibodies, and anti-integrin monoclonal antibodies (2). The TDAG8 agonist BTB09089 has been developed and recently investigated for anti-inflammatory properties. BTB09089 was shown to activate TDAG8 *in vitro,* but provided weak activity *in vivo* according to one study (34). An additional study has shown *in vivo* efficacy of BTB09089 using an ischemic stroke murine disease model (28). Further studies must be done to develop highly efficacious TDAG8 agonists for potential use as IBD therapeutics.

## Acknowledgements

We would like to thank Luke Ashley, Nancy Leffler, Elizabeth Krewson, Lixue Dong, Calvin Justus, Joani Oswald, and Shayan Nik Akhtar for their excellent technical assistance. Additionally, we would like to thank Dr. Owen Witte for providing the TDAG8 KO mice. This study was supported in part by research grants from the National Institutes of Health (1R15DK109484-01, to L.V.Y. and K.L.), Brody Brothers Endowment Fund (#213817, to L.V.Y., H.H., and K.L), and Vidant Cancer Research and Education Fund (to L.V.Y).

---

**Supplementary Fig. 1.**
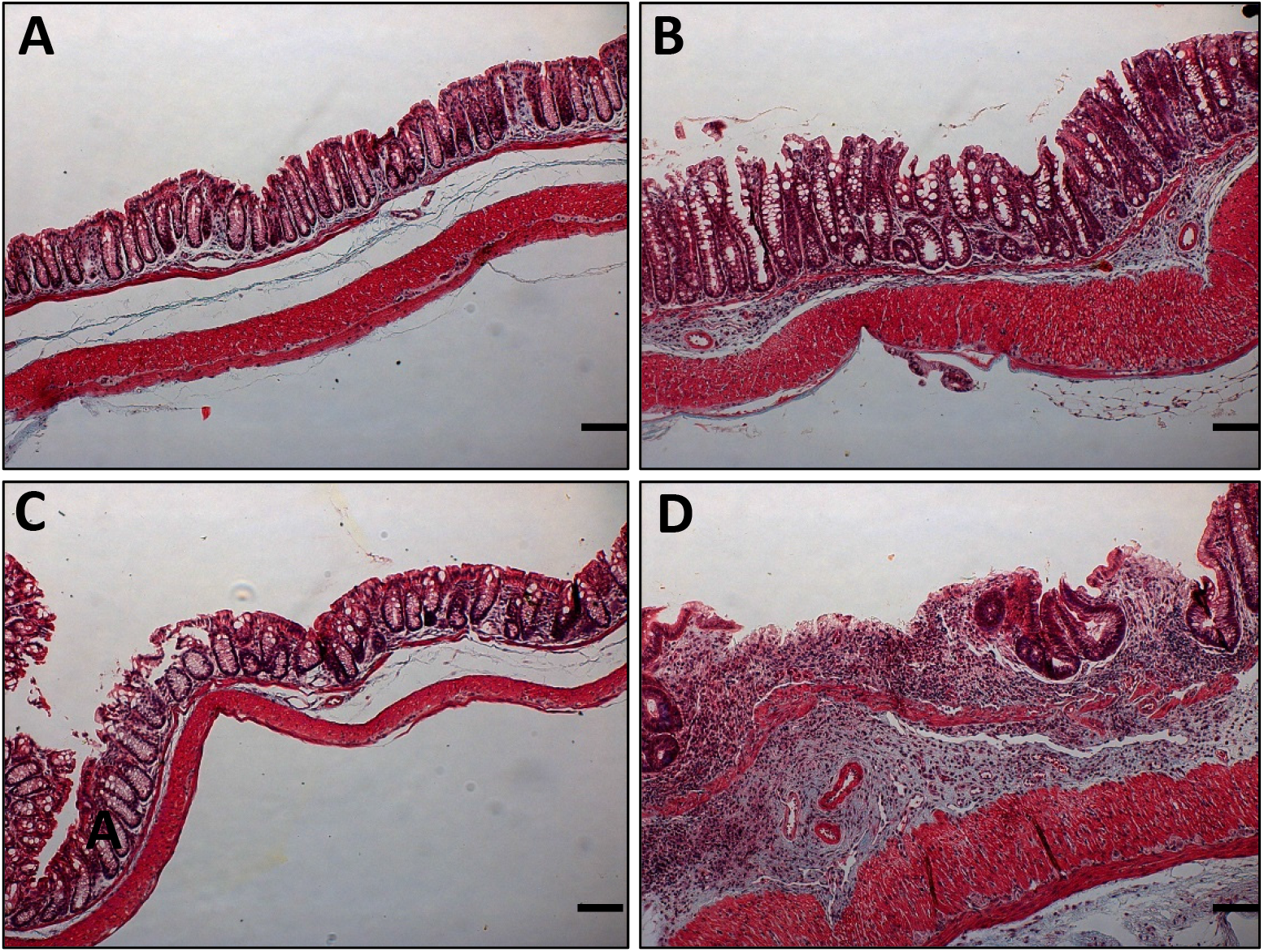
Representative pictures of distal colonic fibrosis. Masson’s trichrome stain indicates DSS-treated TDAG8 KO mice have increased pathological fibrosis compared to WT-DSS mice. (A) WT untreated, (B) WT-DSS, (C) TDAG8 KO untreated, and (D) TDAG8 KO-DSS. Scale bar is 100μm.

**Supplementary Fig. 2.**
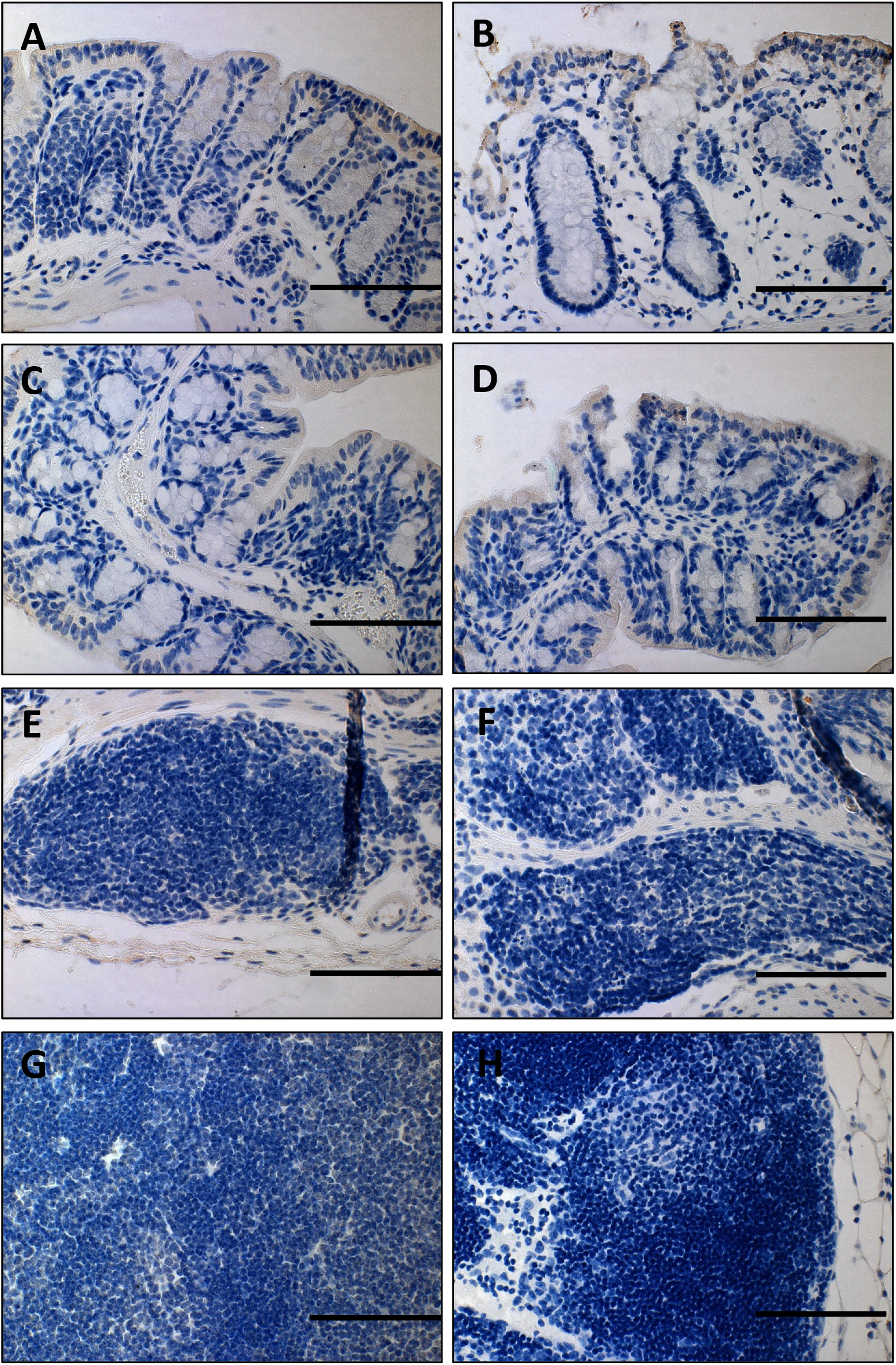
WT negative control for GFP signal in the intestine and intestinal associated lymphoid tissues. No GFP signal could be detected in WT tissues. GFP signal could not be observed in WT-control (A) distal colon mucosa, (C) proximal colon transverse folds, (E) isolated lymphoid follicles (ILFs), and (G) mesenteric lymph nodes (MLNs). GFP signal also could not be detected in WT-DSS (B) intestinal mucosa, (D) transverse folds, (F) ILFs, and (H) MLNs. Minor non-specific staining could be seen on intestinal epithelium and ex-mural connective tissues. Scale bar is 100μm.

## References

1. Al-Mossawi MH, Chen L, Fang H, Ridley A, de Wit J, Yager N, Hammitzsch A, Pulyakhina I, Fairfax BP, Simone D, Yi Y, Bandyopadhyay S, Doig K, Gundle R, Kendrick B, Powrie F, Knight JC, and Bowness P. Unique transcriptome signatures and GM-CSF expression in lymphocytes from patients with spondyloarthritis. Nat Commun 8: 1510, 2017.

2. Chandel S, Prakash A, and Medhi B. Current scenario in inflammatory bowel disease: drug development prospects. Pharmacol Rep 67: 224–229, 2015.

3. Chassaing B, Aitken JD, Malleshappa M, and Vijay-Kumar M. Dextran sulfate sodium (DSS)-induced colitis in mice. Curr Protoc Immunol 104: Unit 15 25, 2014.

4. Chen A, Dong L, Leffler NR, Asch AS, Witte ON, and Yang LV. Activation of GPR4 by acidosis increases endothelial cell adhesion through the cAMP/Epac pathway. PLoS One 6: e27586, 2011.

5. Choi JW, Lee SY, and Choi Y. Identification of a putative G protein-coupled receptor induced during activation-induced apoptosis of T cells. Cell Immunol 168: 78–84, 1996.

6. Ding S, Walton KL, Blue RE, McNaughton K, Magness ST, and Lund PK. Mucosal healing and fibrosis after acute or chronic inflammation in wild type FVB-N mice and C57BL6 procollagen alpha1(I)-promoter-GFP reporter mice. PLoS One 7: e42568, 2012.

7. Dong L, Krewson EA, and Yang LV. Acidosis Activates Endoplasmic Reticulum Stress Pathways through GPR4 in Human Vascular Endothelial Cells. Int J Mol Sci 18, 2017.

8. Dong L, Li Z, Leffler NR, Asch AS, Chi JT, and Yang LV. Acidosis Activation of the Proton-Sensing GPR4 Receptor Stimulates Vascular Endothelial Cell Inflammatory Responses Revealed by Transcriptome Analysis. PLoS One 8: e61991, 2013.

9. Fallingborg J, Christensen LA, Jacobsen BA, and Rasmussen SN. Very low intraluminal colonic pH in patients with active ulcerative colitis. Dig Dis Sci 38: 1989–1993, 1993.

10. Galvez J. Role of Th17 Cells in the Pathogenesis of Human IBD. ISRN Inflamm 2014: 928461, 2014.

11. Gaublomme JT, Yosef N, Lee Y, Gertner RS, Yang LV, Wu C, Pandolfi PP, Mak T, Satija R, Shalek AK, Kuchroo VK, Park H, and Regev A. Single-Cell Genomics Unveils Critical Regulators of Th17 Cell Pathogenicity. Cell 163: 1400–1412, 2015.

12. Hardin M, Cho M, McDonald ML, Beaty T, Ramsdell J, Bhatt S, van Beek EJ, Make BJ, Crapo JD, Silverman EK, and Hersh CP. The clinical and genetic features of COPD-asthma overlap syndrome. Eur Respir J, 2014.

13. He XD, Tobo M, Mogi C, Nakakura T, Komachi M, Murata N, Takano M, Tomura H, Sato K, and Okajima F. Involvement of proton-sensing receptor TDAG8 in the anti-inflammatory actions of dexamethasone in peritoneal macrophages. Biochem Biophys Res Commun 415: 627–631, 2011.

14. Hussman JP, Beecham AH, Schmidt M, Martin ER, McCauley JL, Vance JM, Haines JL, and Pericak-Vance MA. GWAS analysis implicates NF-kappaB-mediated induction of inflammatory T cells in multiple sclerosis. Genes Immun 17: 305–312, 2016.

15. Ishii S, Kihara Y, and Shimizu T. Identification of T cell death-associated gene 8 (TDAG8) as a novel acid sensing G-protein-coupled receptor. J Biol Chem 280: 9083–9087, 2005.

16. Jin Y, Sato K, Tobo A, Mogi C, Tobo M, Murata N, Ishii S, Im DS, and Okajima F. Inhibition of interleukin-1beta production by extracellular acidification through the TDAG8/cAMP pathway in mouse microglia. J Neurochem 129: 683–695, 2014.

17. Jostins L, Ripke S, Weersma RK, Duerr RH, McGovern DP, Hui KY, Lee JC, Schumm LP, Sharma Y, Anderson CA, Essers J, Mitrovic M, Ning K, Cleynen I, Theatre E, Spain SL, Raychaudhuri S, Goyette P, Wei Z, Abraham C, Achkar JP, Ahmad T, Amininejad L, Ananthakrishnan AN, Andersen V, Andrews JM, Baidoo L, Balschun T, Bampton PA, Bitton A, Boucher G, Brand S, Buning C, Cohain A, Cichon S, D’Amato M, De Jong D, Devaney KL, Dubinsky M, Edwards C, Ellinghaus D, Ferguson LR, Franchimont D, Fransen K, Gearry R, Georges M, Gieger C, Glas J, Haritunians T, Hart A, Hawkey C, Hedl M, Hu X, Karlsen TH, Kupcinskas L, Kugathasan S, Latiano A, Laukens D, Lawrance IC, Lees CW, Louis E, Mahy G, Mansfield J, Morgan AR, Mowat C, Newman W, Palmieri O, Ponsioen CY, Potocnik U, Prescott NJ, Regueiro M, Rotter JI, Russell RK, Sanderson JD, Sans M, Satsangi J, Schreiber S, Simms LA, Sventoraityte J, Targan SR, Taylor KD, Tremelling M, Verspaget HW, De Vos M, Wijmenga C, Wilson DC, Winkelmann J, Xavier RJ, Zeissig S, Zhang B, Zhang CK, Zhao H, Silverberg MS, Annese V, Hakonarson H, Brant SR, Radford-Smith G, Mathew CG, Rioux JD, Schadt EE, et al. Host-microbe interactions have shaped the genetic architecture of inflammatory bowel disease. Nature 491: 119–124, 2012.

18. Justus CR, Dong L, and Yang LV. Acidic tumor microenvironment and pH-sensing G protein-coupled receptors. Front Physiol 4: 354, 2013.

19. Justus CR, Sanderlin EJ, Dong L, Sun T, Chi JT, Lertpiriyapong K, and Yang LV. Contextual tumor suppressor function of T cell death-associated gene 8 (TDAG8) in hematological malignancies. J Transl Med 15: 204, 2017.

20. Karnes JH, Cronin RM, Rollin J, Teumer A, Pouplard C, Shaffer CM, Blanquicett C, Bowton EA, Cowan JD, Mosley JD, Van Driest SL, Weeke PE, Wells QS, Bakchoul T, Denny JC, Greinacher A, Gruel Y, and Roden DM. A genome-wide association study of heparin-induced thrombocytopenia using an electronic medical record. Thromb Haemost 113: 772–781, 2015.

21. Kim JJ, Shajib MS, Manocha MM, and Khan WI. Investigating intestinal inflammation in DSS-induced model of IBD. J Vis Exp, 2012.

22. Kottyan LC, Collier AR, Cao KH, Niese KA, Hedgebeth M, Radu CG, Witte ON, Khurana Hershey GK, Rothenberg ME, and Zimmermann N. Eosinophil viability is increased by acidic pH in a cAMP- and GPR65-dependent manner. Blood 114: 2774–2782, 2009.

23. Kyaw H, Zeng Z, Su K, Fan P, Shell BK, Carter KC, and Li Y. Cloning, characterization, and mapping of human homolog of mouse T-cell death-associated gene. DNA Cell Biol 17: 493–500, 1998.

24. Lardner A. The effects of extracellular pH on immune function. J Leukoc Biol 69: 522–530, 2001.

25. Lassen KG, McKenzie CI, Mari M, Murano T, Begun J, Baxt LA, Goel G, Villablanca EJ, Kuo SY, Huang H, Macia L, Bhan AK, Batten M, Daly MJ, Reggiori F, Mackay CR, and Xavier RJ. Genetic Coding Variant in GPR65 Alters Lysosomal pH and Links Lysosomal Dysfunction with Colitis Risk. Immunity, 2016.

26. Latella G, Di Gregorio J, Flati V, Rieder F, and Lawrance IC. Mechanisms of initiation and progression of intestinal fibrosis in IBD. Scand J Gastroenterol 50: 53–65, 2015.

27. Lund PK and Zuniga CC. Intestinal fibrosis in human and experimental inflammatory bowel disease. Curr Opin Gastroenterol 17: 318–323, 2001.

28. Ma XD, Hang LH, Shao DH, Shu WW, Hu XL, and Luo H. TDAG8 activation attenuates cerebral ischaemia-reperfusion injury via Akt signalling in rats. Exp Neurol 293: 115–123, 2017.

29. Mogi C, Tobo M, Tomura H, Murata N, He XD, Sato K, Kimura T, Ishizuka T, Sasaki T, Sato T, Kihara Y, Ishii S, Harada A, and Okajima F. Involvement of proton-sensing TDAG8 in extracellular acidification-induced inhibition of proinflammatory cytokine production in peritoneal macrophages. J Immunol 182: 3243–3251, 2009.

30. Mosenden R and Tasken K. Cyclic AMP-mediated immune regulation--overview of mechanisms of action in T cells. Cell Signal 23: 1009–1016, 2011.

31. Murata N, Mogi C, Tobo M, Nakakura T, Sato K, Tomura H, and Okajima F. Inhibition of superoxide anion production by extracellular acidification in neutrophils. Cell Immunol 259: 21–26, 2009.

32. Nagasaka A, Mogi C, Ono H, Nishi T, Horii Y, Ohba Y, Sato K, Nakaya M, Okajima F, and Kurose H. The proton-sensing G protein-coupled receptor T-cell death-associated gene 8 (TDAG8) shows cardioprotective effects against myocardial infarction. Sci Rep 7: 7812, 2017.

33. Nugent SG, Kumar D, Rampton DS, and Evans DF. Intestinal luminal pH in inflammatory bowel disease: possible determinants and implications for therapy with aminosalicylates and other drugs. Gut 48: 571–577, 2001.

34. Onozawa Y, Fujita Y, Kuwabara H, Nagasaki M, Komai T, and Oda T. Activation of T cell death-associated gene 8 regulates the cytokine production of T cells and macrophages in vitro. Eur J Pharmacol 683: 325–331, 2012.

35. Onozawa Y, Komai T, and Oda T. Activation of T cell death-associated gene 8 attenuates inflammation by negatively regulating the function of inflammatory cells. Eur J Pharmacol 654: 315–319, 2011.

36. Radu CG, Cheng D, Nijagal A, Riedinger M, McLaughlin J, Yang LV, Johnson J, and Witte ON. Normal immune development and glucocorticoid-induced thymocyte apoptosis in mice deficient for the T-cell death-associated gene 8 receptor. Mol Cell Biol 26: 668–677, 2006.

37. Raker VK, Becker C, and Steinbrink K. The cAMP Pathway as Therapeutic Target in Autoimmune and Inflammatory Diseases. Front Immunol 7: 123, 2016.

38. Rieder F and Fiocchi C. Intestinal fibrosis in inflammatory bowel disease - Current knowledge and future perspectives. J Crohns Colitis 2: 279–290, 2008.

39. Rohle D, Popovici-Muller J, Palaskas N, Turcan S, Grommes C, Campos C, Tsoi J, Clark O, Oldrini B, Komisopoulou E, Kunii K, Pedraza A, Schalm S, Silverman L, Miller A, Wang F, Yang H, Chen Y, Kernytsky A, Rosenblum MK, Liu W, Biller SA, Su SM, Brennan CW, Chan TA, Graeber TG, Yen KE, and Mellinghoff IK. An inhibitor of mutant IDH1 delays growth and promotes differentiation of glioma cells. Science 340: 626–630, 2013.

40. Sanderlin EJ, Justus CR, Krewson EA, and Yang LV. Emerging roles for the pH-sensing G protein-coupled receptors in response to acidotic stress. Cell Health Cytoskelet 7: 99–109, 2015.

41. Sanderlin EJ, Leffler NR, Lertpiriyapong K, Cai Q, Hong H, Bakthavatchalu V, Fox JG, Oswald JZ, Justus CR, Krewson EA, O’Rourke D, and Yang LV. GPR4 deficiency alleviates intestinal inflammation in a mouse model of acute experimental colitis. Biochim Biophys Acta 1863: 569–584, 2017.

42. Tcymbarevich I, Richards SM, Russo G, Kuhn-Georgijevic J, Cosin-Roger J, Baebler K, Lang S, Bengs S, Atrott K, Bettoni C, Gruber S, Frey-Wagner I, Scharl M, Misselwitz B, Wagner CA, Seuwen K, Rogler G, Ruiz PA, Spalinger M, and de Valliere C. Lack of the pH-sensing Receptor TDAG8 [GPR65] in Macrophages Plays a Detrimental Role in Murine Models of Inflammatory Bowel Disease. J Crohns Colitis, 2018.

43. Tsurumaki H, Mogi C, Aoki-Saito H, Tobo M, Kamide Y, Yatomi M, Sato K, Dobashi K, Ishizuka T, Hisada T, Yamada M, and Okajima F. Protective Role of Proton-Sensing TDAG8 in Lipopolysaccharide-Induced Acute Lung Injury. Int J Mol Sci 16: 28931–28942, 2015.

44. Wang JQ, Kon J, Mogi C, Tobo M, Damirin A, Sato K, Komachi M, Malchinkhuu E, Murata N, Kimura T, Kuwabara A, Wakamatsu K, Koizumi H, Uede T, Tsujimoto G, Kurose H, Sato T, Harada A, Misawa N, Tomura H, and Okajima F. TDAG8 is a proton-sensing and psychosine-sensitive G-protein-coupled receptor. J Biol Chem 279: 45626–45633, 2004.

45. Xavier RJ and Podolsky DK. Unravelling the pathogenesis of inflammatory bowel disease. Nature 448: 427–434, 2007.

